# Pixelated microfluidics for drug screening on tumour spheroids and ex vivo microdissected primary tissue

**DOI:** 10.1101/2022.10.07.511162

**Authors:** Dina Dorrigiv, Pierre-Alexandre Goyette, Amélie St-Georges-Robillard, Anne-Marie Mes-Masson, Thomas Gervais

## Abstract

Anti-cancer drugs have the lowest success rate of approval in drug development programs. Thus, preclinical assays that closely predict the clinical responses to drugs are of utmost importance in both clinical oncology and pharmaceutical research. 3D tumour models preserve the tumoural architecture and are cost-, labour-, and time-efficient. However, the short-term longevity, limited throughput, and limitations to live imaging of these models have so far driven researchers towards simpler, less realistic tumour models such as monolayer cell cultures. Here, we present a static open-space microfluidic drug screening platform that enables the formation, culture, and multiplexed delivery of several reagents to various 3D tumour models, namely cancer cell line spheroids and ex vivo primary tumour fragments. Our platform utilizes an open-space microfluidic technology, a pixelated chemical display, which creates fluidic “pixels” of biochemical reagents that stream over tumour models in a contact-free fashion. Up to 9 different treatment conditions can be tested over 144 samples in a single experiment. We provide a proof-of-concept application by staining fixed and live tumour models with multiple cellular dyes. Furthermore, we demonstrate that the various responses of the tumour models to biological stimuli can be assessed using the proposed drug screening platform. The platform is amenable to various 3D tumour models, such as tumour organoids. Upscaling of the microfluidic platform to larger areas can lead to higher throughputs, and thus will have a significant impact on developing treatments for cancer.

## Introduction

A major impediment to cancer treatment is predicting the response of patients to anti-cancer drugs as they have an extremely low clinical approval rate in drug development programs. [1, 2] Improving preclinical models to predict the response of patients to treatments can improve drug precision and effectiveness, spare patients from exposure to unnecessary toxicities, accelerate the drug development process, and ultimately reduce healthcare costs. [3],[4] Various predictive preclinical tumour models are available to researchers. Preclinical tumour models include 2D monolayer cultures of cancer cells, 3D tumour models such as cancer cell line spheroids, tumour organoids, ex vivo cultured tumour fragments, and in vivo models, from the simplest to the most complex, respectively. Monolayer cell cultures are easy to replicate but lack the 3D tumour structure and the interactions between the cancer cells and the tumour microenvironment. [5] In vivo models are the gold standard of preclinical models, but their production is time-consuming and labour intensive. They may also fail to predict the clinical efficacy of drugs due to species differences. [6] 3D tumour models can bridge the gap between 2D and in vivo models: unlike 2D monolayers, 3D tumour models mimic the tumoral architecture and are human-derived and easier to work with than in vivo models. [7–12] Three main groups of 3D tumour models exist. In order of increasing complexity and in vivo relevance, they are cell line spheroids, tumour organoids, and ex vivo cultured tumour explants. They can be selected according to the purpose and requirements of a given study. [11] [13] Drawbacks of the various 3D tumour models include limitations of live imaging and interfacing with histopathology, and the generally low throughput and low viability of tissue, especially for ex vivo tumour explants. The most advanced live imaging methods, such as confocal and multiphoton microscopy, are well known to have severe limitations in 3D biology, notably their limited light penetration depth in live tissue and cost. [14] In addition, the universally recognized standard for primary tissue-based clinical decision-making is histopathology [i.e., the practice of preserving tumour tissues in paraffin or a freezing medium and dissecting them into thin (5-10 μm) slices]. [15] To overcome the limitation of live imaging and increase clinical relevance, in particular taking into account routine clinical pathology, it would be advantageous for 3D tumour models to be compatible with standard histopathology practice. Various techniques have been developed for culture and drug screening on 3D tumour models and preparing them for histopathological analyses. The most conventional technique is culturing tumour models in plastic well plates. Samples are subjected to reagents manually or using robotic liquid handlers in wells. Sample manipulation using pipettes, whether manually or using pipetting robots, imposes risks such as aspirating or shearing the sample while changing the medium. In addition, removing the samples out of the wells for further histopathology processing is tedious. Moreover, plastic well plates are not optimal for preserving the viability and metabolic activity of fragile 3D tumour models, such as ex vivo tumour explants. Our group has previously studied the ex vivo survival of tumour tissue explants and has shown that tumour tissue slices cultured in non-perfused well plates start to die in two days due to insufficient oxygen supply. [16] Microfluidics can palliate this problem by introducing chips for high throughput processing of micron-sized tumour models. [5, 17] The drawback of most microfluidic chips is that they require sample entrapment in closed microchannels, and are not amenable to surface-based work environments such as Petri dishes. [18] Perifusion-based microfluidic devices have been developed to preserve the viability of larger tissue explants over a longer time. [19, 20] It is important to differentiate between perfusion and perifusion. [21] Tumour tissues are dense structures with permeabilities that are orders of magnitude below the permeability of flow channels. [22, 23] Creating convective flow inside tumour models (i.e., perfusion) is not feasible unless flow around them is prevented. In most cases, when samples are small (< 1 mm), perifusion is sufficient to avoid any form of starvation, anoxia and necrosis in tissue. [24] Perifusion-based devices, while presenting a technical breakthrough, have extremely limited throughputs. [25, 26]

New culture platforms that can improve survival and high throughput drug screening on 3D tumour models, while remaining fully compatible with gold standard tissue analysis, are promising avenues to improve pre-clinical drug testing. In this article, we present a platform that bridges the concepts behind well plates and perifusion-based microfluidics. The platform uses open-space microfluidic laminar flow confinement to stream reagents within self-contained fluidic pixels in which a large number of various 3D tumour models can be placed. The pixels reagent content can be modulated over time with specific frequencies. Open-space microfluidic systems are channel-free and contact-free fluidic processors that deliver reagents directly over the sample. [27] Pioneering open-space microfluidic systems have been used for various purposes including single cell analysis, [28–30] perifusion-based culture of brain slices, [31] localized immunohistochemistry, [32] and imaging mass spectrometry. [33] The open-space microfluidics system here, which we call the Pixelated Chemical Display (PCD), has been used for various processes over flat 2D surfaces, such as immunoassays. [34] Here, for the first time, we have utilized the PCD for multiplexed reagent screening over 3D tumour models. To this end, we investigated the stability of the PCD when working over a large number of 3D biological samples deposited in a custom-built microwells array. To provide proof of concept evidence for the applicability of the platform across a whole spectrum of 3D tumour models, we first worked with the simplest 3D tumour models, spheroids, and later with the most complex 3D models, ex vivo tumour explants. Using sequences of cellular dyes, we performed crosstalk-free tissue staining on both 3D tumour models. We have also adapted our previously published paraffin-embedding lithography to transfer all samples simultaneously to a paraffin or optimal cutting temperature (OCT) compound block while preserving their spatial orientation. Finally, we investigate the feasibility of using this method to study signalling pathways and cell fate in microdissected tumour tissues. The opportunities and challenges of the method are discussed with respect to competing methodologies such as robotic liquid handlers and closed microfluidic chips.

## Results and discussion

### Design and fabrication of the pixelated chemical display drug screening platform

The PCD operates based on the hydrodynamic confinement of a stream of fluid in another miscible fluid through recirculation. [18, 35] It comprises a blunt tip with multiple apertures and is installed in close vicinity of an immersed substrate. The PCD and the immersed substrate form a Hele-Shaw cell, a quasi-2D flow that can be precisely computed using potential flow theory. [36, 37] During its operation over the substrate, fluid streams are expelled through the injection apertures and re-collected through the aspiration apertures. As a result of the convective recirculation, fluid streams leaving the PCD form well-defined patterns over the substrate without mixing. [38] Fluidic “pixels” are created when a fluid stream injected above the surface is confined by neighbouring identical fluid streams, forming a repeating flow unit with translational symmetries. [39] By modulating the design of the PCDs and the injection and aspiration flow rates, different sizes, numbers, and patterns of fluidic pixels can be achieved. Our group has previously demonstrated theoretically and experimentally the operation of up to 144 fluidic pixels (12 × 12) and demonstrated that the number of active pixels and their reagent content can be modulated without altering the stability of the system. [35],[39] Based on our previous findings, we adapted a 9-pixel PCD for tissue culture and drug screening. Each pixel is 36 mm^2^ (6 × 6 mm^2^) such that the resulting array fits within a paraffin cassette for later embedding (Figure 1 a,b). For this work, each group of 3 pixels was connected to the same reagent flask to create experimental triplicates. Three different conditions were tested in each experiment. We designed and micromachined a microwell array to keep the tumour models in place at the PCD interface during the experiment. The microwell array features 9 groups of 16 microwells for a total of 144 microwells on a polymethylmethacrylate (PMMA) slab (Figure 1 b). Each microwell group is covered by an independent fluidic pixel when the PCD is aligned over the microwell array. The PMMA slab also features a flat surface on the side of the microwell array to safely install the PCD and test its operation prior to biological experiments over tumour models (Figure 1 b). 3D printed holder assembly parts (Figure 1 c) were fabricated to securely hold the PCD and the PMMA slab together, to stabilize the PCD, and ensure its alignment over the microwell array (Figure 1 d).

**Figure 1.**
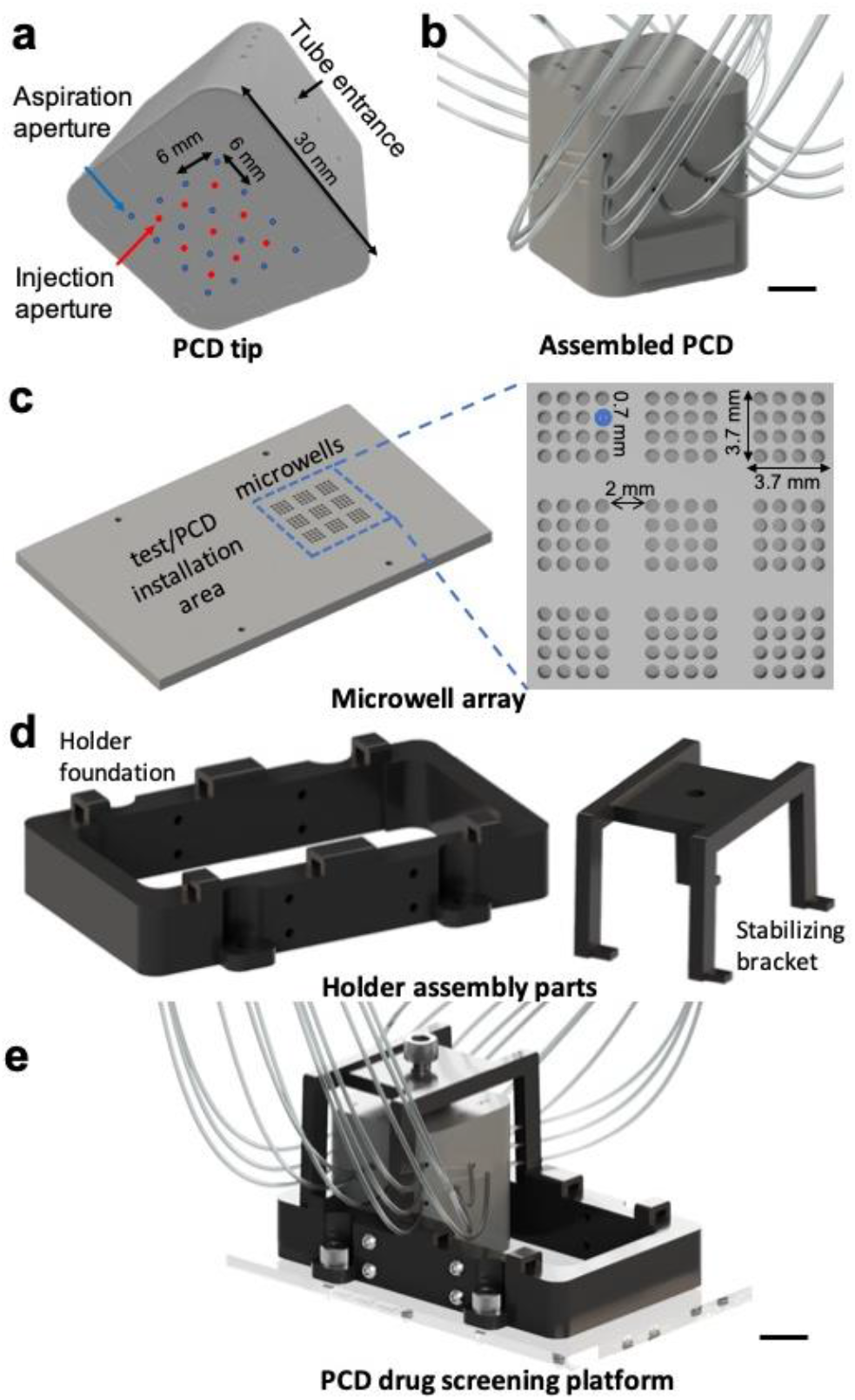

PCD components. a) PCD tip and b) tubes connected to the PCD, c) micromachined microwell array featuring 144 microwells, holder assembly parts: d) holder foundation (left) and a bracket to stabilize the PCD once installed over the microwells (right), and e) schematic of the fully assembled PCD drug screening platform. Scale bar = 1 cm.

### Pressure pump-operated fluidic lines

Syringe pumps were previously used to operate PCDs [34, 39] as they offer precise and simple control, and are commonly used in microfluidic systems. [40] However, syringe pumps, even high precision ones, have relatively high minimum working flow rates, and require frequent recharging of reagents in syringes. [41] To avoid these limitations, we used pressure pumps in this work. Pressure pumps enable pressurizing of a wide range of flask sizes (from microliters to litres capacity) and thus allow for longer-run experiments. More importantly, a single pressure pump can be used to pressurize several reagent flasks, whereas each syringe pump is dedicated to a single syringe. Different flow rates can be achieved in different fluidic lines pressurized by one pump by controlling the hydraulic resistance of the tubes (i.e., by using different sizes of tubing). In this work, two pressure pumps were used to operate the PCD: one for the injection groups and one for the aspiration. Similar to our previous works, we used 3D-printed manifolds to deliver fluids from one pump into all the pixels sharing the same reagents. [39] Tubes connecting the manifolds to the PCD were used as precision hydraulic resistors to match the flow rate from all apertures. Four precision flowmeters were installed on the fluidic lines to measure the flow rate for the four injection and aspiration groups. A closed-loop control system with a feedback loop control (a.k.a., proportional– integral–derivative [PID] controller) was developed to control the pressure-driven flows. The PID controller estimates the deviation between the target and the measured injection flow rates and regulates the pressure to reduce deviations in real time. Moreover, we added medical three-way stopcock valves on the fluidic lines to enable on-demand reagent switching between the various reagent flasks (e.g., priming reagents such as ethanol and isopropanol, culture medium, biochemicals, and cellular dyes). Switch valves enable us to add or remove reagent flasks without interrupting the system (Supplementary Figure S1). We refer to the PCD, microwell array, pumps, and fluidic lines complex as the PCD drug screening platform.

### Finite element simulations

Our group has previously studied the mass transport, stability, and reconfigurability of PCDs using 2D convection-diffusion finite-element methods and has demonstrated that stochastic errors such as minor pressure, flowrate changes, or clogging of one aperture do not impact the PCD’s operation. [39] Here, we conducted numerical simulations to gain insight into the quality, crosstalk, and stability of fluidic pixels when the PCD is working over 3D structures, such as tumour models. We added a secondary set of simulations to predict the convective-diffusive transport of diluted species in microwells and inside tumour models. We used experimental geometrical and operational parameters: flow rates were selected based on the minimum flow rate that we could achieve with the pressure pumps to have sharp and stable pixels while minimizing reagent consumption. To visualize the pixel formation and crosstalk, we modelled a PCD that functioned in a chequerboard pattern: five injection apertures inject a concentrated solution, and four injection apertures inject a zero-concentration solution (Figure 2 a). Simulation results suggest that the presence of microwells and tumour models does not disturb the pixel shapes, similarly to working over flat impermeable surfaces [Figure 2 a(i)]: stable crosstalk-free pixels are formed over the microwell array [Figure 2 a(ii)]. Moreover, we investigated the impact of the tumour model positioning in microwells on fluidic pixels. We modelled a scenario in which random tumour models were partially sticking out of the wells and touched the PCD. We observed that regardless of the tumour model positioning in the well, the pixels are stable (Figure 2 b). We then evaluated the shear stress induced on the cells by the flow and observed that the maximum shear stress imposed by the PCD is 0.0021 Pa, which is 500 times less than the physiologically safe shear stress regime for sensitive cells (^~^1 Pa). [42] By visualizing velocity fields, we observed that there is no convective flow inside the tumour models [Figure 2 c], showing the diffusion dominant transfer of species inside tumour models. Next, we measured the amount of time required to reach a constant concentration of injected species inside the tumour models with zero initial concentration. Simulation results predict that the system reaches a steady state in less than 20 minutes. The time to reach this state was set as the transition time in the experiments; upon the change of a reagent flask, reagents were streamed for 20 minutes before starting the experiment countdown. Overall, our simulations predict that the PCD provides excellent control over the fluidic pixels, and locally perifuses the tumour models. Finally, to model the spheroid formation assay (i.e., the static culture of 500 μm-tumour models in the microwell array), we used passive diffusion and Michaelis-Menten kinetics parameters for oxygen and glucose consumption by cancer cell lines. [24] The numerical model predicts that spheroids have access to sufficient levels of oxygen and glucose over 24 hours (i.e., the typical medium refreshment interval) (Figure 2 d).

**Figure 2.**
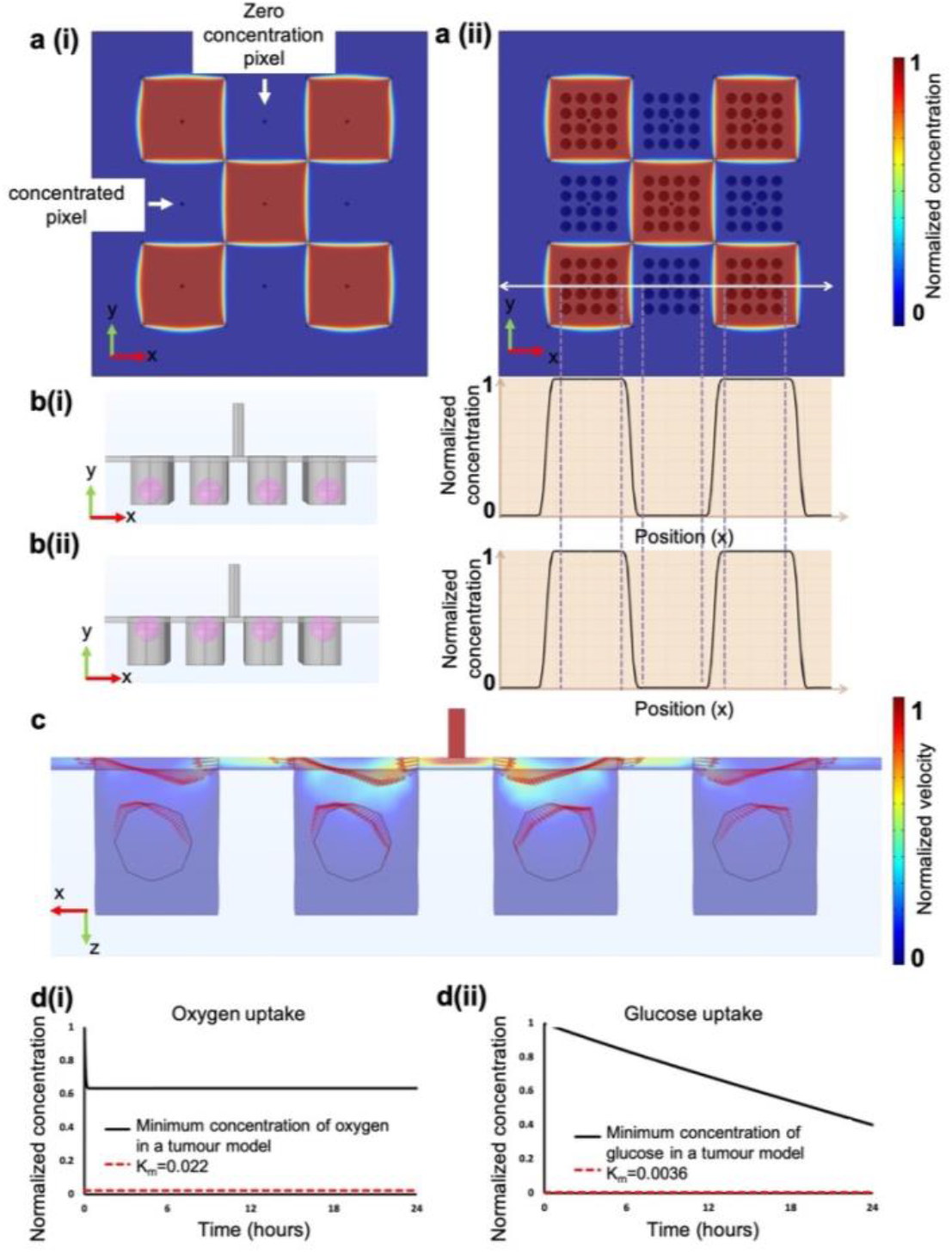

Numerical simulations of the PCD operation over tumour models. a) Fluidic pixels formed over a flat surface a(i) are comparable to those formed over the microwell array a(ii); b) The positioning of the MDTs or spheroids in the microwells does not impact the operation of the PCD; c) arrow plots to visualize the distribution of velocity field in the numerical model suggest the lack of free flow inside tumour models; d(i) Oxygen and d(ii) glucose consumption profile in tumour models cultured in the microwell array without perifusion. Tissues of up to 500 μm in diameter can survive in microwells for over 24 hours, as the oxygen and glucose concentration stays above typical K_m_ values (normalized) for cancer cells.

### High-throughput formation of cancer cell line spheroids is possible in the microwell array

Cancer cell line spheroids are self-formed spherical cell aggregates formed from one or more cancer cell lines. [43] Spheroids are the simplest and most often used 3D tumour models. The commonly used technique to form spheroids is to seed a high-density suspension of cells on a non-adherent surface. Spheroids will form if the cell-cell adhesion forces are greater than the cell-surface adhesion forces. [44] To meet our claim about the amenability of PCD to work with different tumour models, we optimized the surface modification technique and cell seeding densities based on previous findings to form spheroids directly in the microwell array. [45–47] We further formed spheroids from squamous cell carcinoma and colorectal cancer cell lines to test the practicality of the approach. Figure 3 shows that we can form uniform spheroids in the microwell array in 48 hours. Spheroids were later subjected to the PCD for dynamic cellular staining.

**Figure 3.**
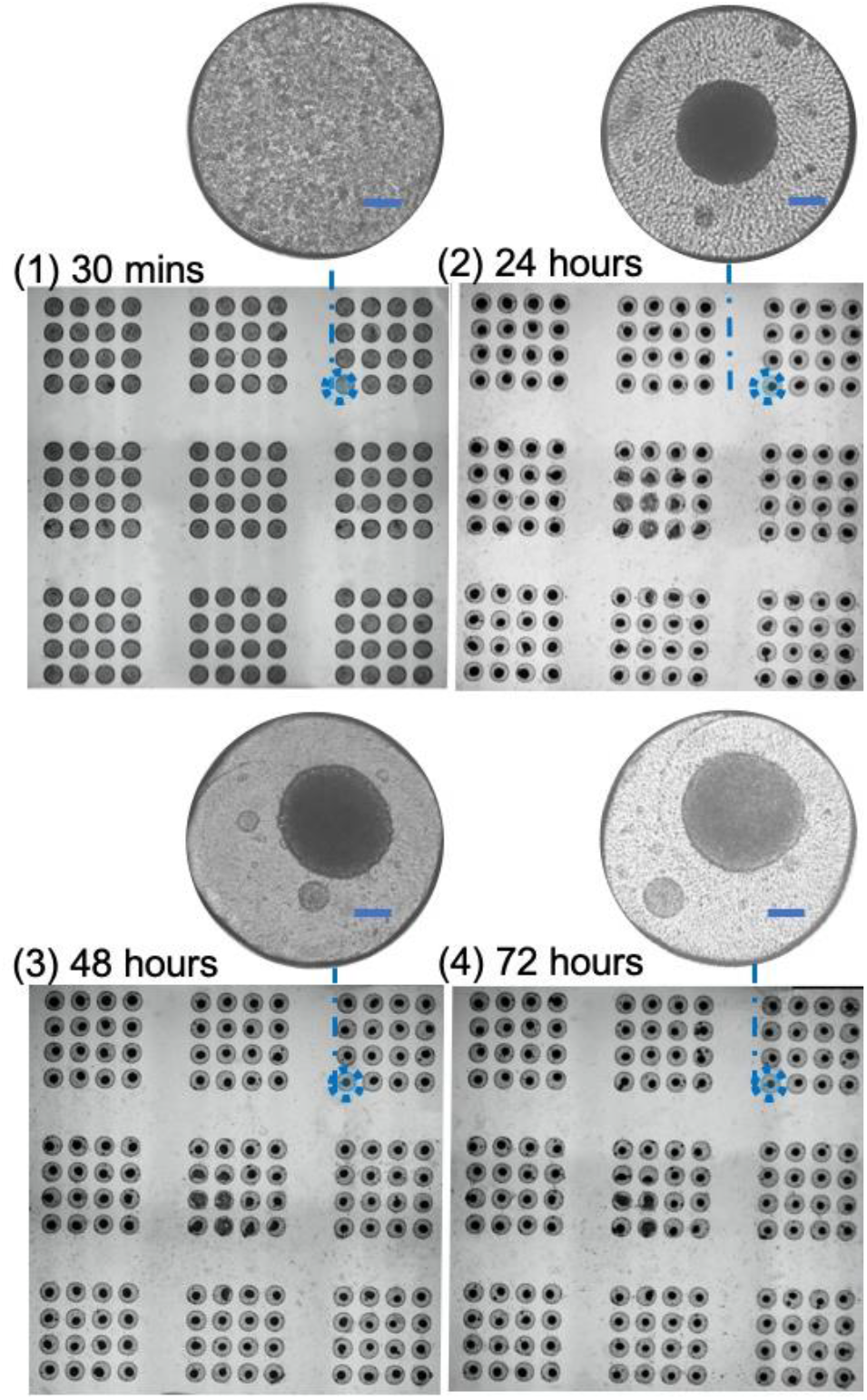

Formation of uniform and compact spheroids of colon cancer cell line HCT-116 in the microwell array over time. Scale bar = 100 μm.

### The PCD drug screening platform enables dynamic multiplexed staining of spheroids

To test the performance, stability, and precision of the drug screening platform, we used it to stain spheroids formed in the microwell array. Two phases of reagent streaming over spheroids were used to test the stability of the PCD over subsequent changes of reagents, and to verify its potential for dynamic reagent screening. The PCD streamed culture medium for 20 minutes over the microwell array containing spheroids. Subsequently and without interrupting the system, the culture medium was replaced by three cellular dyes that were streamed in the 9 pixels of the PCD for 2 hours. We then switched the reagent flasks, subjecting spheroids to a second dye. The second part of reagent streaming went on for 3 hours, and cellular dyes were swapped with PBS 1X to purge the dyes (Figure 4 a). Spheroids were imaged using fluorescence microscopy. The results show crosstalk-free staining of spheroids with the colours of interest (Figure 4 b). We further assessed the fluorescent intensity (FI) per unit area of spheroids for different channels and demonstrated that spheroids subjected to a dye for 3 hours have a higher FI than spheroids subjected to the same dye for 2 hours (Figure 4 c).

**Figure 4.**
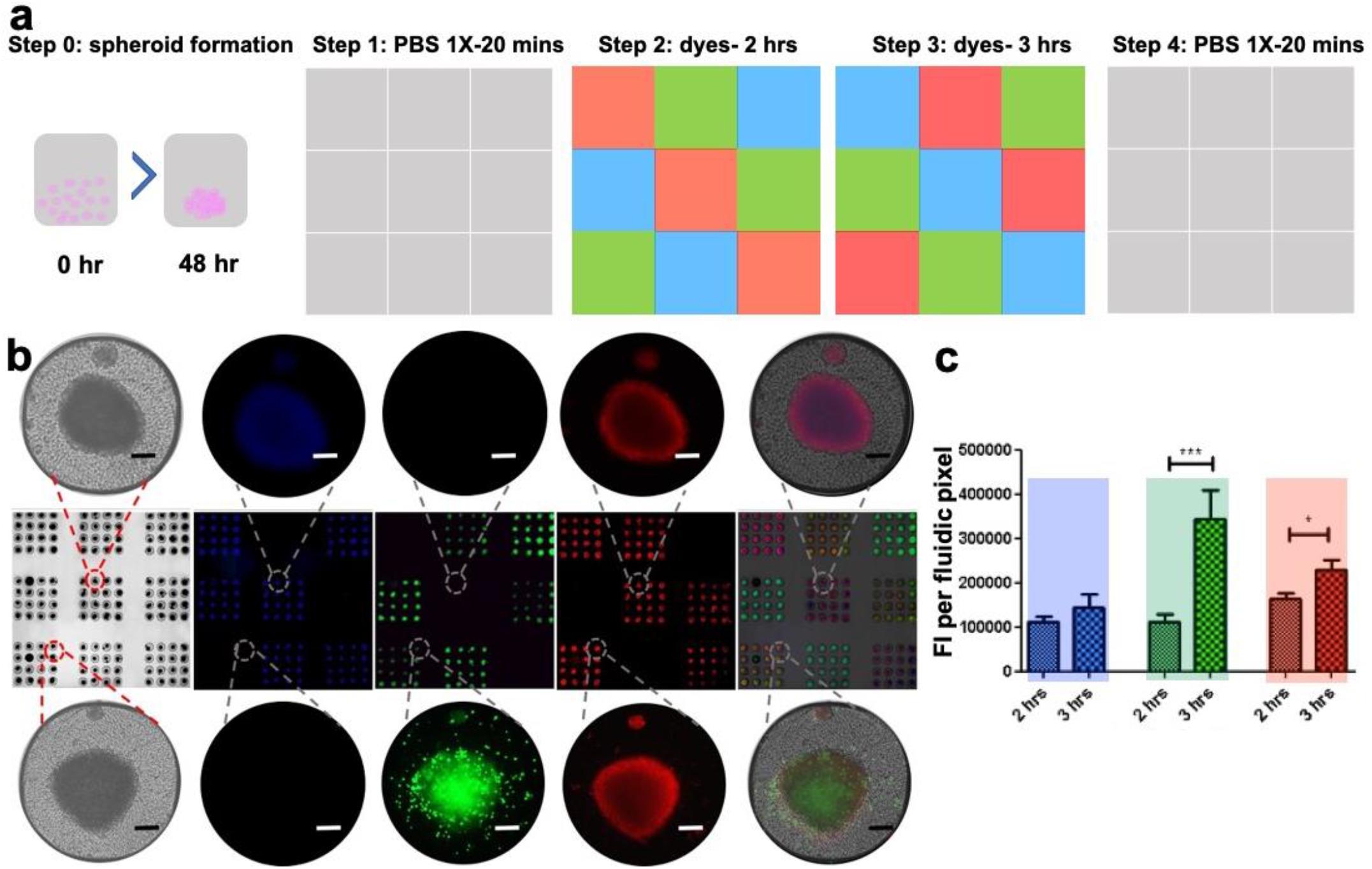

Crosstalk-free multiplexed staining of tumour models using the PCD. The PCD is used to stain HCT-116 spheroids 48 hours after cell seeding. Spheroids are exposed spheroids are subjected to 3 different cellular dyes streaming at the 9 pixels of the PCD for two hours. Then, the reagents were switched so that a different dye was streamed at each pixel for 3 hours. Spheroids were imaged after rinsing out the dyes using an inverted fluorescent microscope. The staining protocol (a), micrograph of stained spheroids (b), and quantification of Fluorescent Intensity (FI) of spheroids for each channel (c). Longer incubation with fluorophores results in higher fluorescent emission of spheroids. Blue: Hoechst, green: Celltracker™ Green, red: Celltracker™ Red. Scale bar = 100 μm.

### The PCD drug screening platform can process fragile *ex vivo* tumour tissue explants

Ex vivo tumour tissue explants provide an excellent tumour model since they are readily available from biopsy or surgery, do not require disintegration, and mirror the individual’s tumour features including the histological and gene expression profiles. [17, 48] However, they are the least frequently used 3D tumour model due to the frailty of the tissue and limited throughput. Our group has devised a methodology in which tumour tissues are dissected into sub-millimetre-sized fragments and cultivated on microfluidic chips. [49] The micro-dissected tumour tissue (MDT) methodology yields many MDTs from a small primary tissue and maximizes tissue viability. [17] Its applications are restricted by the fact that limitations are imposed due to its closed microfluidic chips. To free the system from these limitations, we investigated the application of the PCD drug screening platform to MDTs. Fresh or formalin-fixed MDTs produced from xenografted tumour tissue were deposited in the microwell array and subjected to the PCD. The PCD streamed cellular dyes over the MDT. Images of whole MDTs captured using the fluorescent microscope showed crosstalk-free staining of MDTs with the intended colours, similar to what was observed for spheroids (Supplementary Figure S2).

### Tumour tissue microarray

Fluorescence microscopy captured the mean fluorescence emission of tumour model structures but did not allow us to examine the distribution of reagents throughout the tumour models. To demonstrate this capability in our system, we developed a methodology to take tumour models out of the microwells and directly embed them in a freezing medium or paraffin. It is also essential to keep the arrangement and orientation of tumour models as they were in the microwells to be able to correlate them with the treatment conditions (in each fluidic pixel) that they have been exposed to. For this, we adapted a technique previously described by Jones and Calabresi [50] to first embed the tumour models in a hydrogel in their microwells. Then, the hydrogel block containing the tumour models is de-moulded from the microwell array and re-embedded in the OCT compound. The OCT blocks were then sectioned into 5 μm-thick slices, and slices were used for further histopathological staining and analysis (Figure 5). It is noteworthy that this protocol makes the PCD drug screening platform compatible with standard histopathology practice. Supplementary Figure S2 shows cryosections of MDTs that were stained using the PCD for different durations and that underwent the agarose and OCT embedding protocol.

**Figure 5.**
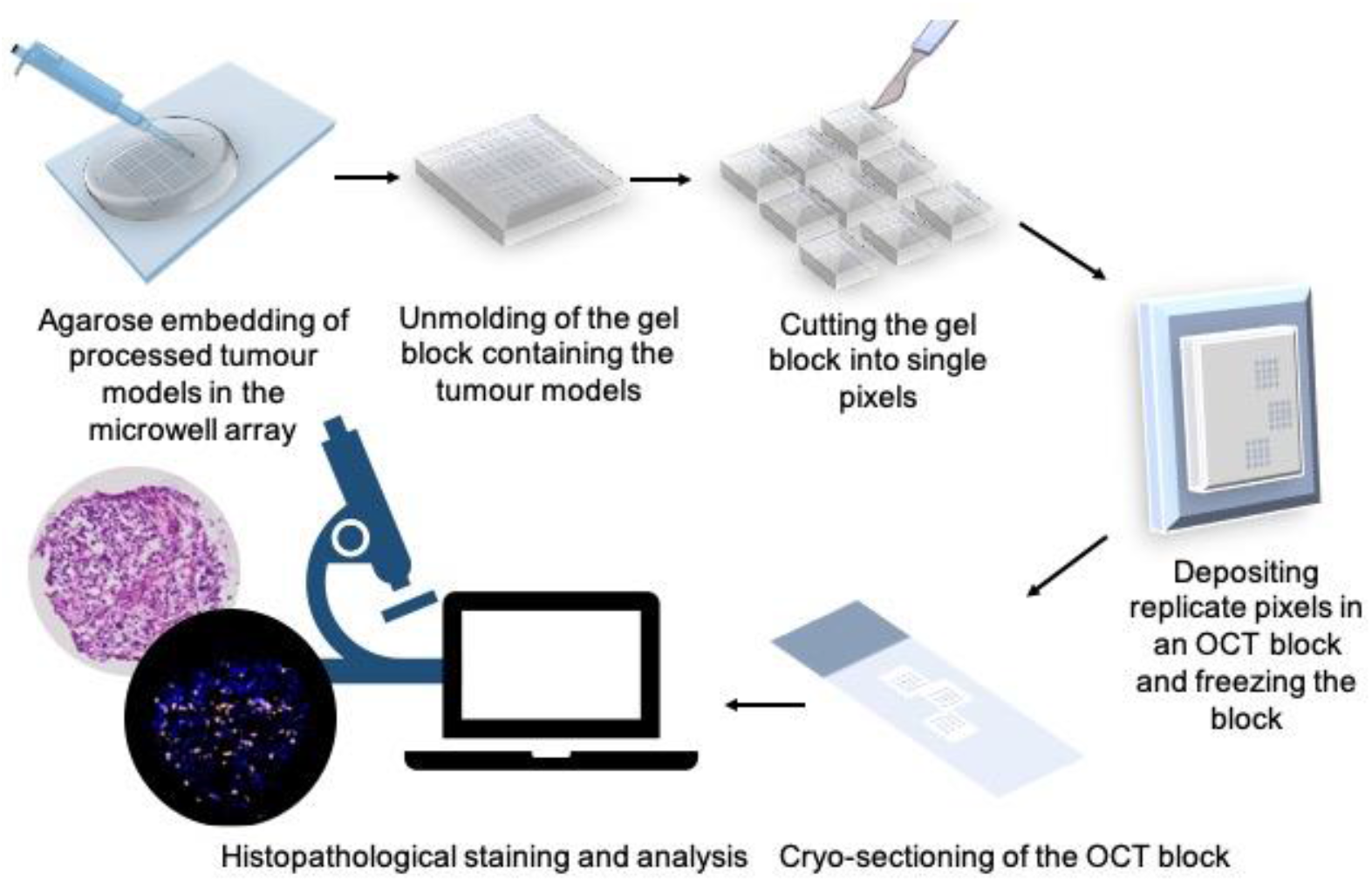

The protocol developed to remove the tumour models from microwells while preserving their address for further histopathology analyses. Tumour models are embedded in agarose and removed from the microwell array. Microwell groups exposed to different fluidic pixels are separated, and tumour models that have been subjected to the same treatment condition are regrouped in an OCT block. OCT blocks are sectioned to 5 μm sections to visualize the tissue core and for further immunostaining.

### The PCD drug screening platform enables the tracking of biological responses in tumour models

After demonstrating that the PCD platform is capable of forming crosstalk-free fluidic pixels over various tumour models, we sought to examine the ability of the technology to follow various biological responses in tumour models. For this, we assessed the response of Nuclear factor kappa B (NF-κB) transcription factors in MDTs to a cytokine (tumour necrosis factor [TNF]) stimulation. The NF-κB transcription factor is reported to play a role in tumour angiogenesis and invasiveness and is a possible target to improve the clinical diagnosis and prognosis. [51] NF-κB resides in the cytoplasm of every cell and is translocated to the nucleus when activated by various stimuli such as cytokines, viruses, and free radicals. [52] TNF is a proinflammatory cytokine that is known to activate NF-κB. [53] Real-time monitoring of nuclear translocation of Nf-κB proteins in previous studies revealed a rapid increase in the nuclear signal of sub-units of NF-κB that peaks a few minutes after the exposure, followed by a decline in the nuclear signal of proteins. [54, 55] With this in mind, we used the PCD to expose MDTs produced from cell line xenografts to TNF for 0, 30 minutes, or 240 minutes by progressively switching on TNF delivery in certain pixels by replacing neutral culture medium with a TNF solution. We evaluated the nuclear signal of p65, an NF-κB subunit, by immunofluorescence (IF) staining. As expected, the quantification of the IF staining showed an increase in the nuclear signal of p65 in MDTs that have been subjected to TNF stimulus for 30 minutes compared to the control group. The p65 nuclear signal dropped in the MDTs that were treated for 240 minutes (Figure 6). This is also expected since p65 translocation is known to be reversible. [53] To further validate the results, we performed parallel experiments on MDTs produced from the same xenograft that were cultured on chips. The on-chip MDT treatment experiment is a repeat of a protocol previously published by our laboratory for other cell line xenograft MDTs. [17] Similar responses were also observed in MDTs on chips (Figure 6). 2D cell cultures of the same cell line treated with TNF showed similar results (Supplementary Figure S3), substantiating the results seen in MDTs. The results showcase that the PCD drug screening platform can reflect the response of tumour samples to biological stimuli.

**Figure 6.**
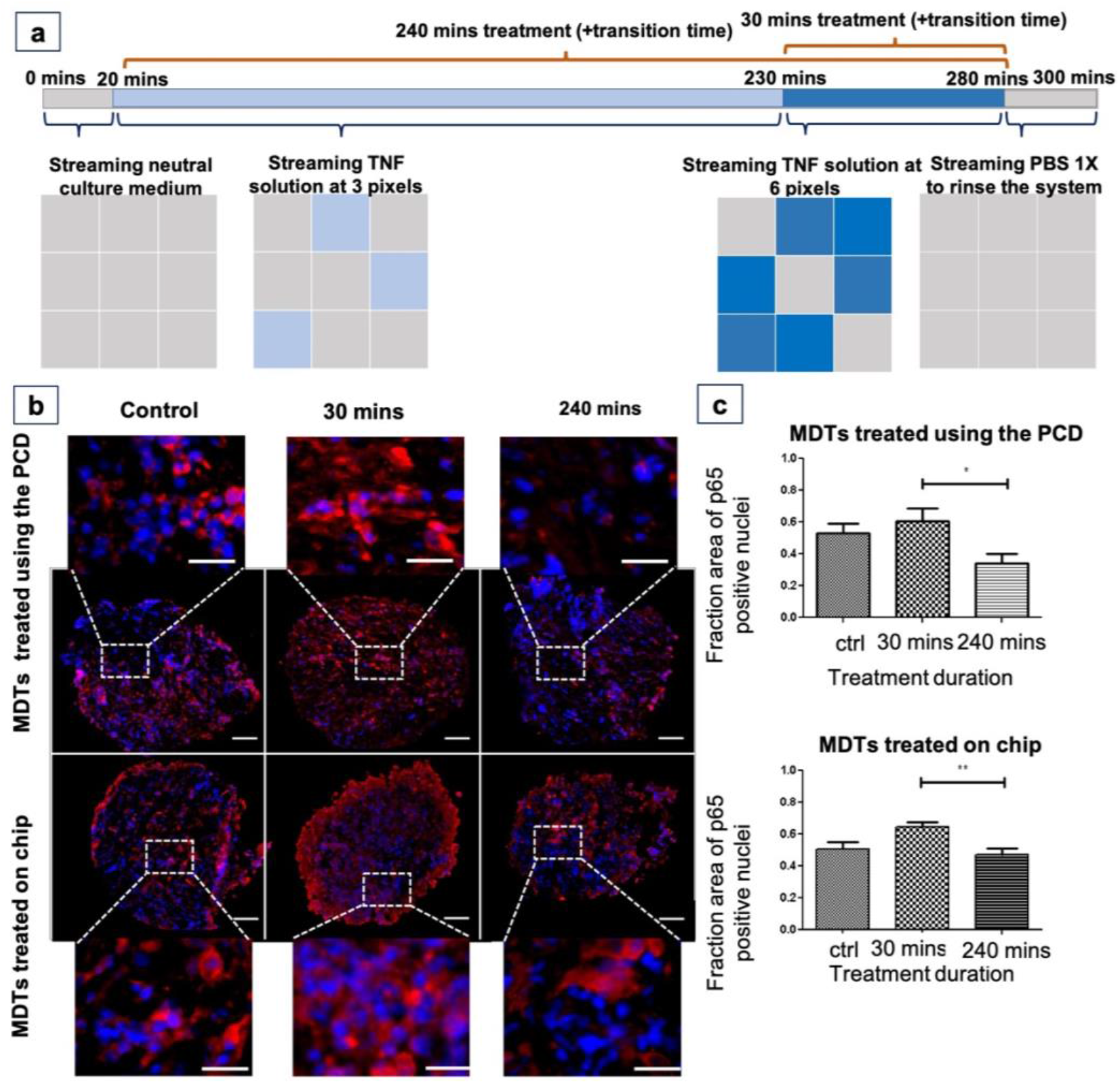

The PCD drug screening platform can recreate the on-chip response of tumour models to a cytokine. Xenograft cell line MDTs (TOV 21G) were treated with a TNF solution for different durations using the PCD and on-chip (a), and the change in the nuclear translocation of p65 was quantified (b-c). Red: p65 and blue: DAPI. Scale bar = 20 μm for zoomed-in MDTs, and 100 μm for whole MDTs. N = 3.

## Material and methods

### Design and fabrication of parts

We followed our previously published methodology to design and fabricate PCDs, manifolds, and the holder assembly. [39] Briefly, all parts except PCDs and manifolds were designed in Fusion 360 (Autodesk Inc., CA, USA) software. PCDs and manifolds were designed using script-assisted CAD in Catia V5 (Dassault Systèmes, France) previously developed by our group. [34] The parts were 3D printed using a stereolithography 3D printer (PICO2 HD, Asiga, Australia). The resin used was Pro3dure GR-1 black (P3GR1BLK-1L, Pro3dure medical GmbH, Iserlohn, Germany). After the printing, the excess resin was cleaned by sonication of the parts in an isopropanol bath, then post-cured by UV exposure (Flash UV Curing Chamber, Asiga). To assemble the PCD, 1/16” (RK-06419-01, Masterflex Tygon, Cole-Parmer, Quebec, Canada) and 1/32” (RK-06420-01, Masterflex Tygon, Cole-Parmer) tubing was plugged and glued using a UV-sensitive resin. Polycarbonate three-way stopcock valves (RK-30600-02, Cole-Parmer) were installed on the fluidic lines. Glue and screws were used to assemble the parts of the holder assembly. The microwell array was micromachined using an MDX-40A milling machine (Roland DGA, Irvine, CA, USA) on a 1/8” PMMA slab (8560K239, McMaster-Carr, Elmhurst, IL, USA). The holder assembly was fixed over the PMMA slab. Polydimethylsiloxane (PDMS; Sylgard® 184 silicone elastomer kit, Dow Corning, Midland, MI, USA) was used to seal the holder assembly-PMMA slab interface and form a liquid-tight environment inside the holder assembly. Flow rates were controlled using AF1 microfluidic pressure pumps and MFS4 microfluidic flow sensors (Elveflow, Paris, France). Microfluidic chips for drug testing on MDTs were fabricated using our previously published protocol. [56]

### System operation

The operation of the system was controlled using custom Python (Python software foundation) and LabView codes (National Instrument, Austin, TX, USA). A flow rate of 0.5 μl/s per aperture was used for all reagent streaming over live tissue. The aspiration to injection flowrate ratio was kept at 1.4. To prepare the system, isopropanol was first streamed at 1 μL/s per aperture in all of the injection and aspiration tubes for at least 15 minutes to wet, prime, and sterilize the fluidic lines. The system was further infused for 5 minutes with PBS 1X (PBS 10X; 3072318, Wisent Inc., Saint-Bruno-de-Montarville, Canada) in all lines to purge isopropanol. Next, without pausing the pumps, the PCD was installed in the holder assembly over the immersed flat surface beside the microwell array. The experimental flow rates for injection and aspiration were administered, and the PCD was gently slid over the microwell array. PBS 1X for formalin fixed tissue, or neutral culture medium for live tissue, was administered for 20 minutes to rinse the microwells. Next, the reagents of interest were put in place. For formalin-fixed tissue staining experiments, Sytox™ Green (S7020, Thermo Fisher Scientific, Waltham, MA, USA), Nuclear Mask Red (H10326, Thermo Fisher Scientific), and DAPI (D1306, Thermo Fisher Scientific) were used. 10 mM solution of DAPI in PBS 1X was prepared and aliquoted. Dye solutions were diluted at a 1:500 ratio in PBS 1X. For live tissue staining experiments, Celltracker™ Green (CMFDA; C2925, Thermo Fisher Scientific), Celltracker™ Red (CMRA; C34551, Thermo Fisher Scientific), and Hoechst (Hoechst 33342, H3570, Thermo Fisher Scientific) were used. Dye solutions were diluted at a 1:500 ratio in the culture medium. At the end of the experiment, PBS 1X was injected for at least 10 minutes to rinse the fluidic lines and tumour models from the reagents. Experiments were performed on a microscope stage and at room temperature. Injection reagent flasks were put in a water bath at 40 °C. Heating the reagent helps to prevent bubble formation in the fluidic lines and over the microwell array.

### Finite element methodology

We used COMSOL Multiphysics© software v.5.6 (COMSOL Inc, Burlington, MA, USA) to simulate the convection and diffusion of species under the PCD and within tissue models. Passive diffusion of oxygen and glucose in the static culture of tumour models in between the medium changes was also modelled for the spheroid formation assay. The geometry of the model was drawn using built-in COMSOL drawing tools. The dimensions of the model can be found in Supplementary Table S1. All simulations were conducted at a constant biological temperature (37°C). We used a time-dependent solver to model the PCD tumour model system and the spheroid formation assay. For the PCD tumour model system, Fick’s second law of diffusion and Navier-Stokes equation for laminar flow were applied using the “transport of diluted species in porous medium” module. The injection apertures were considered as inflows (i.e., source), injecting species with two interchangeably varying concentrations (i.e., concentrated or zero concentration solutions): starting from the top right side of the PCD tip, every other pixel received the concentrated solution, resulting in 5 pixels streaming the concentrated solution and 4 pixels streaming the zero-concentration solution. The aspiration apertures were considered as outflows (i.e., sink). The aspiration to injection flow rate ratio was optimized to yield sharp pixels at the experimental flow rates and was kept constant throughout the simulation. The operational parameters of the model can be found in Supplementary Table S2. All the liquid compartments of the model had the physical properties (i.e., density and viscosity) of water at 37 °C. The porosity and hydraulic permeability are extremely low for the tumour model compartment. [22, 23] With this in mind, we assumed the tumour models non-porous and used the diffusion coefficient of glucose in water to model the transport of concentrated solution in the tissue models. For the spheroid formation assay, the transport of diluted species module was used to model the passive uptake of glucose and oxygen by tumour models. We first simulated oxygen transfer within the tumour models in the spheroid formation assay. We considered a constant oxygen concentration at the medium-air interface over the microwell array. PMMA is not gas-permeable, thus we imposed no-flux (Neumann) boundary conditions at the bottom and walls of the microwells. For glucose, we assumed continuity boundary conditions at the medium-tumour model interface. We used Michaelis–Menten (MM) kinetics to model cancer cells’ glucose and oxygen consumption rates in the spheroid formation assay. The average Michaelis–Menten uptake kinetics found in the literature [24, 49, 57] imply high consumption rates in the abundance of nutrients and decreased consumption rates when nutrients are depleted. The Michaelis–Menten constants refer to concentration thresholds, below which the normal cell metabolism is impacted. [58] We evaluated the minimum concentration of oxygen and glucose in the core of tissues of 500 μm in diameter. Tissue uptake and diffusion parameters are provided in Supplementary Table S2.

### Cancer cell lines xenograft tumour production

A human carcinoma cell line derived from an ovarian cancer tumour TOV21G (RRID:CVCL_3613) was used to produce mouse xenografts. Ovarian cancer cells were grown as monolayers (2D culture) in OSE medium (316-030-CL, Wisent Inc.) supplemented with 10% fetal bovine serum (FBS; Gibco™, Thermo Fisher Scientific), 55 mg/L gentamicin (Gibco™, Thermo Fisher Scientific) and 0.6 mg/L amphotericin B (Gibco™, Thermo Fisher Scientific). After reaching confluency, cells were detached with 0.25% trypsin-EDTA solution (Life Technologies, California, USA), and cell suspensions (1 000 000 cells) were mixed with Matrigel (BD Biosciences, Franklin Lakes, NJ, USA) at a 1:1 ratio and subcutaneously injected into the flank of immunodeficient NOD.Cg-Rag1tm1Mom Il2rgtm1Wjl/SzJ female (Charles River, Wilmington, MA, USA). Xenograft tumours were harvested once they reached a volume between 1 500 and 2 000 mm^3^. All animal procedures were performed in accordance with the Guidelines for the Care and Use of Laboratory Animals of the CRCHUM and approved by the Animal Ethics Committee (the Comité Institutionnel de Protection des Animaux).

### MDT production from cell line xenograft tumours

We used our previously published method [17, 56] for the production of MDTs. Briefly, a tissue chopper (McIlwain, Ted Pella, Redding, CA, USA) was used to cut the xenograft into 350 μm-thick tissue slices. Tissue slices were kept in Hank’s Balanced Saline Solution (HBSS, 311-516-CL, Wisent Inc.) supplemented with serum and antibiotics. Tissue slices were further punched into MDTs using a 500 μm diameter tissue punch (Zivic Instruments, Pittsburgh, PA, USA) and kept in HBSS supplemented with antibiotics.

### Microwell preparation and MDT loading in the microwell array

Similar to PDMS devices [49], the microwell arrays were wetted and rendered hydrophilic by plasma treatment and rinsed with 100% ethanol. They were then sterilized by soaking in 70 % ethanol for 15 minutes and prepared by incubation with a triblock copolymer (Pluronic^®^ F-108, Sigma-Aldrich, St. Louis, MO, USA) overnight (at least 16 hours) at 37 °C in a 5% CO_2_ incubator. The microwell arrays were then rinsed with PBS 1X three times to purge the Pluronic^®^ F-108 solution. We adapted the previously published method of our laboratory to load the MDTs in the microwells. [17, 56] Briefly, the overlay liquid over the microwells was removed. 16 MDTs were picked using a 20 μL pipette and emptied over a microwell group. MDTs were diverted towards empty microwells using the pipette tip where they would fall in the microwells. In the case of more than one MDT falling in a microwell, the extra MDTs were pipetted out of the well and transferred to empty wells. This process was repeated for all 9 microwell groups.

### Spheroid formation assay

We used a human squamous cell carcinoma FaDu (RRID: CVCL_1218) and a human colon cancer cell line HCT-116 (RRID: CVCL_0291) for spheroid formation experiments. Cells were grown as monolayers (2D culture) in Dulbecco’s Modified Eagle Medium (DMEM; 11965118, Gibco™, Thermo Fisher Scientific) supplemented with serum and antibiotics. After reaching confluency, cells were detached and cell suspensions of 2 000 000 cells in 1 ml of culture medium were prepared. 400 μL of cell suspension were seeded over each microwell array, and the cell suspension was replenished three times to exchange the liquid in the microwells with the cell suspension. 15 minutes after the cell seeding, the cell suspension over the microwell array was removed by drawing 400 μL of the cell suspension and adding 400 μL of medium to remove the floating cells over the microwell array. Medium was changed every 24 hours by adding 400 μL of fresh medium near one corner of the microwell array, removing 400 μL from the opposite corner, and repeating the process three times.

### OCT embedding protocol

Following fresh tissue experiments, some tumour models underwent formalin fixation in the microwell arrays. 400 μL of formalin was added near one corner of the microwell array and removed from the opposite corner and repeated three times. The tumour models were incubated in formalin for 40 minutes and formalin was rinsed by three washes with PBS 1X. For agarose embedding, an 8% solution of agarose (Ultrapure™ Low Melting Point Agarose; 16520100, Thermo Fisher Scientific) in PBS 1% was prepared by dissolving 8 g of agarose in 100 ml PBS 1X and microwaving the solution for 80 seconds (4 cycles of 20 seconds) or until agarose powder was completely dissolved. The agarose in PBS solution was then cooled down to 62 °C. 400 μL of agarose solution was discharged and removed three times over the microwell array using a positive displacement pipette. The microwell array was placed in an oven at 60 °C for 30 minutes to ensure agarose permeates in the microwells and tissues. The microwell arrays were further cooled at 4 °C for 30 minutes, and the agarose layer was peeled off gently. If tumour models were left in the microwells after the removal of the agarose, a needle was used to remove the tumour models from the wells and add them to the agarose tissue array. The agarose block was cut to separate the microwell groups, and microwells subjected to the same treatment condition were placed in the same plastic mold, ensuring that tissues were touching the bottom of the plastic mold. OCT was poured gently over the agarose block to prevent bubble formation. Plastic molds were placed on a flat and levelled surface in dry ice and cooled down for 20 minutes for OCT to solidify. Each OCT block was sliced into 5 μm-thick sections using a cryostat, and each section was placed on a TOMO® hydrophilic adhesion slide (Matsunami, Bellingham, WA, USA).

### Histopathological staining

OCT sections underwent hematoxylin and eosin (H&E) staining as well as IF staining to assess the expression of p65 protein (Anti-NFkB p65 protein; SC-8008, Santa Cruz, Texas, USA) and DAPI in the tumour models. IF staining was performed using the BenchMark XT automated stainer (Ventana Medical System Inc., Tucson, AZ). Antigen retrieval was carried out with Cell Conditioning 1 (#950-123, Ventana Medical System Inc) for 90 minutes for all primary antibodies. Mouse anti-p65 (1:200) antibody was automatically dispensed. The slides were incubated at 37 °C for 60 minutes and secondary antibodies were incubated at room temperature on the bench. We used our laboratory’s protocol to quantify the TNF response in 2D culture. [59] Briefly, cells were seeded onto coverslips at 20 000 cells/well in 24-well plates. After 24 h, cells were incubated with TNF solution for 5 minutes or 2 hours. Cells were fixed with formalin for 15 minutes at room temperature, washed using PBS 1X, permeabilized with 0.25% Triton (Triton™ X-100 solution; 93443, Sigma-Aldrich), and incubated with mouse anti-p65 (1:400) overnight. Primary antibody was detected by incubation with secondary antibody for 60 minutes. Coverslips were mounted onto slides using Prolong Gold® anti-fade reagent with DAPI (14209 S, Life Technologies Inc.). All sections were scanned with a 20x/0.75 NA objective with a resolution of 0.3225 μm (bx61vs, Olympus, Toronto, Ontario).

### Tumour model treatment with TNF

For cytokine stimulation experiments, MDTs were exposed to a neutral culture medium or to a 20 ng/ml of TNF solution (Recombinant TNF alpha human; 300-01A, PeproTech, Thermo Fisher Scientific) in culture medium for either 30 minutes or 240 minutes. The TNF treatment using the PCD lasted 300 minutes. First, the PCD streamed neutral culture medium at every pixel for 20 minutes. Then, we streamed TNF in one group (3 pixels) while the two remaining groups (6 pixels) received a neutral culture medium. At 230 minutes, TNF streaming was started in a second group. For the next 50 minutes, TNF was streaming at 6 pixels, all the while the control group on the same 3 pixels received culture medium. At 280 minutes, we swapped all reagent flasks for PBS 1X, and PBS 1X was streamed for 20 minutes to rinse the tumour models. The PCD was then removed, and the immersion liquid over the microwell array was withdrawn. Tumour models underwent OCT embedding for further histopathology processes. We followed our group’s protocol for MDT treatment on-chip. [17]

### Quantification of immunofluorescent staining

To measure the FI of tumour models stained using the PCD, an open-source image processing software (Fiji) was used. [60] At least 3 spheroids were randomly selected in each fluidic pixel, and the corrected FI per area (subtracting the background FI from tissue FI) was calculated for each fluorescent channel. The average corrected FI per area of the 3 pixels subjected to the same treatment was compared between the 2- and 3-hour incubation time for each channel. To quantify protein expressions using immunofluorescent staining, we used VisiomorphDP software (VisioPharm, Hørsholm, Denmark) [40,41]. Briefly, the tissue core surface area was detected through the DAPI channel. The nuclear signal of p65 was quantified by dividing the surface area of p65-positive nuclei by the total surface area of the nuclei. We used a similar approach for quantifying p65 translocation in 2D culture of the cells. [61]

### Statistical analysis

Statistical analyses were performed in GraphPad Prism version 8.0 (San Diego, CA, USA) using the non-parametric one-way ANOVA Kruskal–Wallis test and post hoc Dunn’s test, because the data were not normally distributed according to the D’Agostino and Pearson omnibus normality test. For TNF treatment experiment, a minimum of 15 MDTs were analyzed for each condition, and experiments were repeated three times (N = 3). All data are reported as the mean ± standard error of the mean (SEM) unless otherwise stated. The reported p-values were generated using a post hoc test (Dunn’s test).

## Conclusions

The need to improve the predictive power of in vitro and ex vivo model systems to maximize the chances of success in clinical trials has made 3D tumour models, such as microdissected tissue and cancer cell line spheroids, attractive in preclinical settings. [62, 63] Thus, it is essential to implement tools and techniques to automate drug screening on 3D tumour model systems and make them compatible with clinical practices. To address this, we introduced a drug screening platform for automated simultaneous streaming of up to 9 reagents on 144 tumour models. We used human cancer cell line xenograft and spheroid models to validate the potential of the PCD drug screening platform for multiplexed and dynamic streaming of biochemicals over tumour models. Microtissues processed in the drug screening platform can directly be transferred to an embedding medium and undergo various endpoint measurements (i.e., immunohistochemistry, immunofluorescence, and H&E). These measurements are standard protocols in clinical and pharmaceutical practices and allow the monitoring of multiple biological pathways. Furthermore, because the platform is amenable to different 3D tumour models, it allows the co-culture of spheroids, organoids, and ex vivo tumor tissue explants. In turn, this enables comparing treatment efficacy on various tumour models. The main drawback of the PCD arises from the continuous streaming of reagents. Even though flow rates are extremely low, streaming over several hours consumes a considerable amount of reagents, and thus limits applications in cases where reagents are extremely expensive (e.g., recombinant protein drugs). However, a highly parallel drug screening assay using the PCD would probably even be worthwhile despite the high reagent consumption. We have reported a low number of large pixels (9 × 6 mm^2^) in this article. However, PCDs of up to 144 × 1 mm^2^ pixels have been produced routinely in our laboratory with successive reagent changes as fast as 1 change per 30 s. This makes the PCD drug screening platform appealing for highly parallel and dynamic assays. The reconfigurable sizes and numbers of pixels along with the fast reagent change will speed up the throughput.

Compared to traditional well plate-based approaches [64] and their downsized microfluidic counterparts such as InSphero GravityTRAP™, [65] idenTx™, [66] and Organoplate® [67], our platform does not require the manual delivery of reagents to microtissues or the use of robotic liquid handlers. This should greatly reduce the time and cost required to perform the experiments. Moreover, the possibility of the direct transfer of tumour models to an embedding medium reduces the potential tissue damage and makes our platform more efficient compared to previous approaches where each individual sample is transferred from well plates to an embedding medium.

## Author contributions

Conceptualization: D.D., P.-A.G., A.-M.M.-M. and T.G.; Data curation, D.D.; Formal analysis, D.D.; Funding acquisition, A.-M.M.-M. and T.G.; Investigation, D.D., A.-M.M.-M. and T.G.; Methodology, D.D., P.-A.G., A.S.R., A.-M.M.-M. and T.G.; Project administration, A.-M.M.-M. and T.G.; Resources: A.-M.M.-M. and T.G.; Supervision, A.-M.M.-M. and T.G.; Validation, D.D., A.S.R., P.-A.G., A.-M.M.-M. and T.G.; Visualization, D.D., and A.S.R.; Writing— original draft, D.D. and T.G.; Writing—review and editing, A.S.R., and T.G. All authors have read and agreed to the published version of the manuscript.

## Acknowledgements

We acknowledge Kim Leclerc-Desaulniers for technical assistance with the animal work. We thank the CRCHUM Microfluidics Core Facility supported by the TransMedTech Institute and its main financial partner, the Canada First Research Excellence Fund and namely Jennifer Kendall-Dupont and Benjamin Péant for performing tissue dissection and useful scientific and technical discussions. We thank Liliane Meunier and Véronique Barrès of the CRCHUM Molecular Pathology Core Facility for performing the OCT block sectioning and slide scanning. We acknowledge Jacqueline Chung for manuscript editing. T.G. acknowledges CMC Microsystems.

## Supplementary material

### Supplementary figures

**Figure S1:**
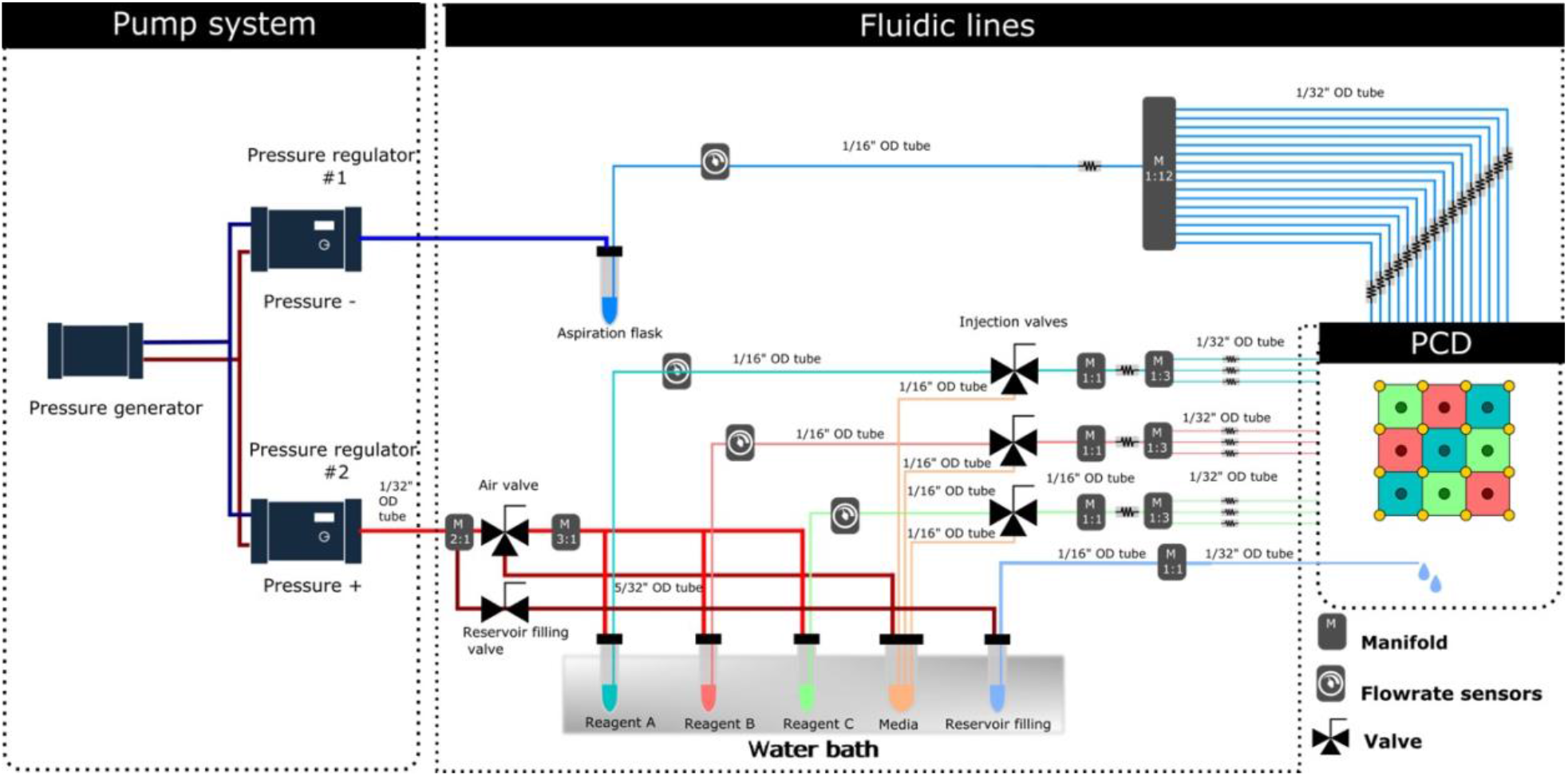
Fluidic connections in the PCD drug screening platform. The use of valves allows for switching between the streaming of various reagents and the flowrate sensors allow for control and validation that the platform is working correctly. OD; outside diameter

**Figure S2:**
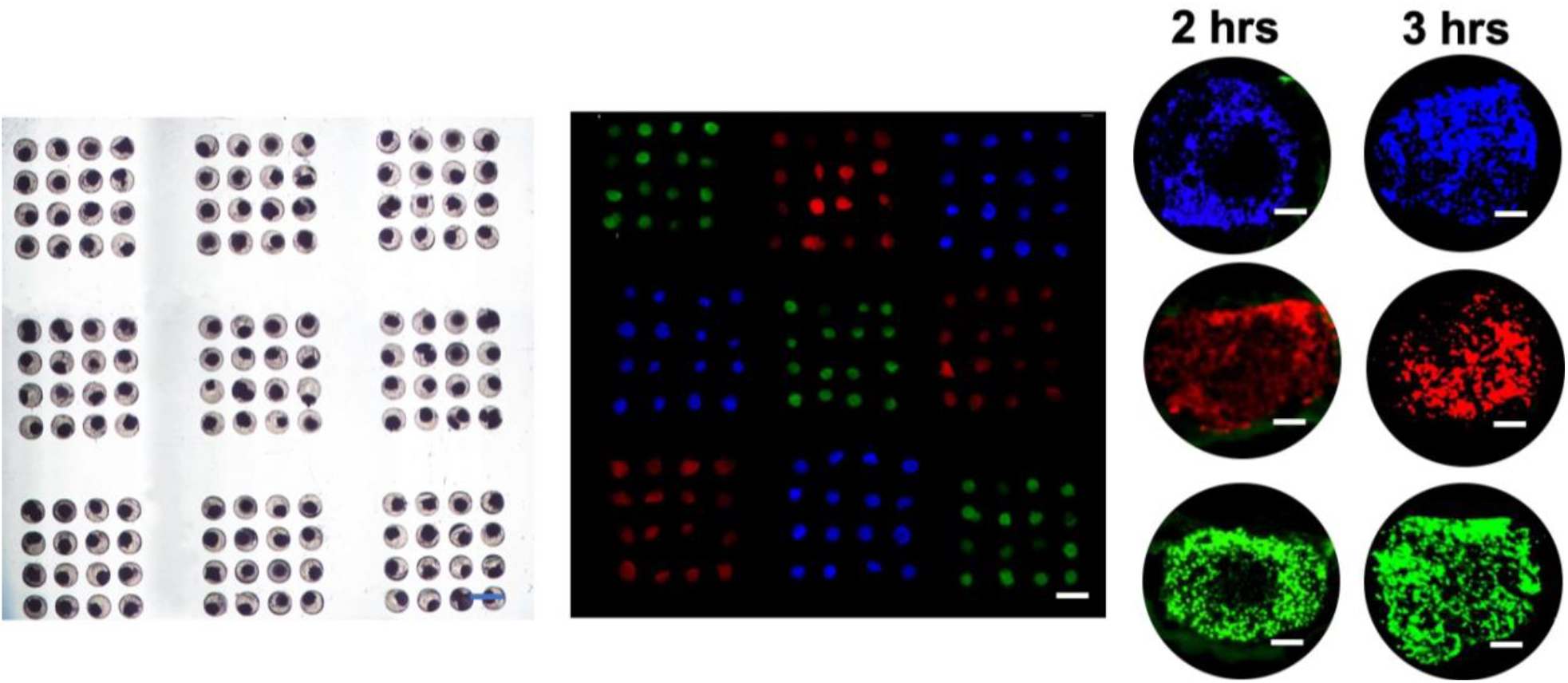
Micrographs of MDTs stained with various cellular dyes using the PCD. Images taken from the tumour model cores that have been treated with cellular dyes for different amounts of time show that core cells are not stained in the shorter treatment durations.

**Figure S3:**
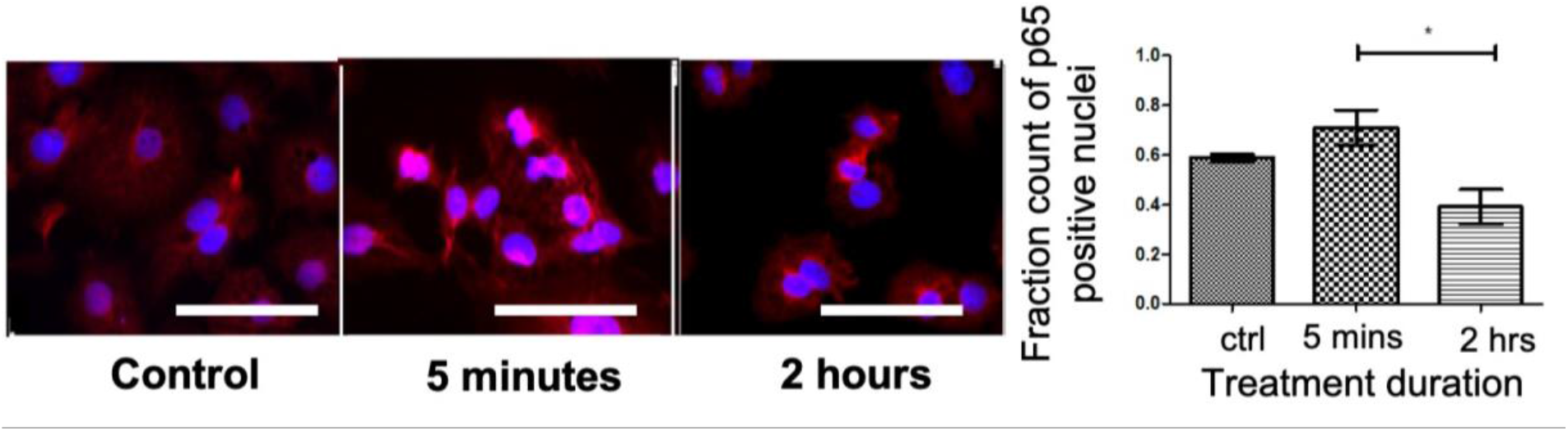
time-dependent treatment of 2D culture of TOV21G cells with TNF shows a response similar to MDTs treated on-chip or using the PCD. This further validates the potential of the PCD drug screening platform to predict the response of 3D tumour models to stimuli.

### Supplementary tables

**Table S1.** dimensions

| Parameter | Value ($\mu\text{m}$) |
| --- | --- |
| **Microwell array** |  |
| well height | 900 |
| well width | 700 |
| well length | 700 |
| Distance between microwells | 1000 |
| **PCD** |  |
| Gap between the PCD and the microwell array | 100 |
| Aperture diameter | 200 |
| Distance between apertures (i.e., pixel size) | 5000 |
| Number of aspiration apertures | 16 |
| Number of injection apertures | 9 |
| **Tissue** |  |
| Tissue diameter | 450 |

**Table S2.** tissue uptake parameters, diffusion properties, the PCD working condition

| Parameter | value |  |  |
| --- | --- | --- | --- |
| Diffusion/Reaction Parameters |  |  |  |
| Diffusion constant of glucose (cm<sup>2</sup>/s) | Tissue | 2.7x10<sup>-6</sup> <sup>1,2</sup> |  |
|  | Medium | 9.27x10<sup>-5</sup> <sup>1</sup> |  |
|  | Agar 5% | Same as water <sup>3,4</sup> |  |
| Diffusion constant of oxygen (cm<sup>2</sup>/s) | Tissue | 1.8x10<sup>-5</sup> |  |
|  | Medium | 2.6x10<sup>-5</sup> |  |
|  | Agar 5% | 2x10<sup>-5</sup> <sup>4</sup> |  |
|  | PDMS | 3.4x10<sup>-5</sup> <sup>5,6</sup> |  |
| Diffusion constant of Carboplatin/Paclitaxel (cm<sup>2</sup>/s) | Could we use glucose properties? <sup>7,8</sup> |  |  |
| Saturation concentration (mM) | Oxygen | Tissue | 1.02 |
|  |  | medium | 0.21 |
|  |  | Agar 5% | 0.21colagen<sup>9</sup> |
|  |  | PDMS | 1.43 |
|  | Glucose | 11 |  |
| Oxygen partition coefficient (relative solubility of oxygen) | PDMS-Medium | 0.15 |  |
|  | Medium-Tissue | 4.8 |  |
| Maximum cellular uptake rate (mM/S) | Oxygen | 2.07 |  |
|  | Glucose | 1.09 |  |
| Michaelis-Menten constant (mM) | Oxygen | 4.63x10<sup>-3</sup> |  |
|  | Glucose | 4x10<sup>-2</sup> |  |
| PCD working conditions |  |  |  |
| Injection pressure (Pa) | 4 |  |  |
| Aspiration pressure (Pa) | 4 |  |  |
| Injection/Aspiration flowrate (nL/s) | 100 |  |  |

